# DNA ligase I fidelity mediates the mutagenic ligation of pol β oxidized nucleotide insertion products and base excision repair intermediates with mismatches

**DOI:** 10.1101/2020.10.30.362251

**Authors:** Pradnya Kamble, Kalen Hall, Mahesh Chandak, Qun Tang, Melike Çağlayan

## Abstract

DNA ligase I (LIG1) completes base excision repair (BER) pathway at the last nick sealing step following DNA polymerase (pol) β gap filling DNA synthesis. We previously reported that pol β 8-oxo-2’-deoxyribonucleoside 5’-triphosphate (8-oxodGTP) insertion confounds LIG1 leading to the formation of ligation failure products with 5’-adenylate (AMP) block. Here, we report the mutagenic ligation of pol β 8-oxodGTP insertion products and an inefficient substrate-product channeling from pol β Watson-Crick like dG:T mismatch insertion to DNA ligation by LIG1 mutant with perturbed fidelity (E346A/E592A) *in vitro*. Moreover, our results revealed that the substrate discrimination of LIG1 for the nicked repair intermediates with preinserted 3’-8-oxodG or mismatches is governed by the mutations at both E346 and E592 residues. Finally, we found that Aprataxin (APTX) and Flap Endonuclease 1 (FEN1), as compensatory DNA-end processing enzymes, can remove 5’-AMP block from the abortive ligation products with 3’-8-oxodG or all possible 12 non-canonical base pairs. These findings contribute to understand the role of LIG1 as an important determinant of faithful BER, and how a multi-protein complex (LIG1, pol β, APTX and FEN1) can coordinate to hinder the formation of mutagenic repair intermediates with damaged or mismatched ends at the downstream steps of the BER pathway.

## INTRODUCTION

Human DNA ligases (DNA ligase I, III, and IV) catalyze the formation of a phosphodiester bond between the 5’-phosphate (P) and 3’-hydroxyl (OH) termini of the DNA intermediate during DNA replication, repair and genetic recombination (1–4). DNA ligation reaction by a DNA ligase includes three chemical sequential steps and requires adenosine triphosphate (ATP) and a divalent metal ion (Mg^2+^) (5). In the first step, ATP is hydrolyzed to produce an adenylate (AMP) that is then covalently linked to the ligase active site lysine, forming the adenylated ligase (LIG-AMP) (6). Next, after binding a nicked DNA substrate, the AMP group is transferred to the 5’-P end of the nick, forming an adenylated DNA intermediate (DNA-AMP) (7). In the final step, DNA ligase catalyzes a nucleophilic attack of 3’-OH in the DNA nick on the adenylated 5’-P to form a phosphodiester bond (8). Despite of the fact that the ligation mechanism is a universally conserved pathway, we still lack an understanding of how DNA ligase recognizes, and processes damaged or mismatched DNA ends.

Successful DNA ligation relies on the formation of a Watson-Crick base pair of the nicked DNA that is formed during prior gap filling DNA synthesis step by a DNA polymerase (9,10). Human DNA polymerases and DNA ligases have been considered as key determinants of genome integrity (11). In our previous studies, we demonstrated the importance of the coordination between DNA polymerase (pol) β and DNA ligase I (LIG1) during the repair of single DNA base lesions through base excision repair (BER) (9,10,12–16). The BER is a critical process for preventing the mutagenic and lethal consequences of DNA damage that arises from endogenous and environmental agents and underlies disease and aging (17,18). The pathway involves a series of sequential enzymatic steps that are tightly coordinated through protein-protein interactions in a process referred to as passing-the-baton or a substrate-product channeling (19–21). This mechanism includes hand off of repair intermediates from gap filling DNA synthesis by pol β to DNA ligation by LIG1 at the downstream steps of the repair pathway (9). However, how deviations in the pol β/LIG1 interplay affect the BER process and genome stability remain largely unknown. In particular, deviations formed due to incorporation of damaged and mismatch nucleotides by pol β could lead to the formation of toxic and mutagenic repair intermediates that can drive genome instability or cell death. For example, endogenous and exogenous oxidative stress can oxidize Guanine in the nucleotide pool (dGTP) and result in the formation of the most abundant form of oxidative DNA damage in nucleotide triphosphate form, 2’-deoxyribonucleoside 5’-triphosphate (8-oxodGTP) (22, 23). The deleterious effect of 8-oxodGTP is mediated through its incorporation into genome by repair or replication DNA polymerases (24). For example, pol β performs mutagenic repair by inserting 8-oxodGTP opposite Adenine at the active site that exhibits a frayed structure with lack of base pairing with a template base after the insertion (25). In our prior studies, we demonstrated that pol β 8-oxodGTP insertion confounds the DNA ligation step of BER pathway and leads to the formation of ligation failure product with 5’-adenylate (AMP) block (12–14). We also reported the effect of pol β mismatch nucleotide insertion on the substrate-product channeling to LIG1 during the final steps of BER and the fidelity of this mechanism in the presence of epigenetically important 5-methylcytosine base modifications (15, 16).

The X-ray crystal structures of the three human DNA ligases in complex with DNA have revealed a conserved three-domain architecture that encircles the nicked DNA, induces partial unwinding and alignment of the 3’- and 5’-DNA ends (26–32). Recently, the crystal structures of LIG1 revealed the important amino acid residues (E346 and E592) that reinforce high fidelity (32). It has been shown that the LIG1 fidelity is mediated by catalytic Mg^2+^-dependent DNA binding that the enzyme employs during the adenyl transfer and nick-sealing steps of ligation reaction. Moreover, the mutations at two glutamic acids residues to alanine (E346A/E592A or EE/AA) leads to the LIG1 with lower fidelity and create an open cavity that accommodates a damaged DNA terminus at the ligase active site (32). However, how such a distinct environment that the staggered LIG1 conformation creates due to the mutations in the important high fidelity sites could affect the ligase substrate discrimination and coordination with pol β during the substrate-product channeling of the repair intermediates with damaged and mismatched DNA ends at the downstream steps of the BER pathway remain entirely undefined.

Aprataxin (APTX), a member of the histidine triad (HIT) superfamily, hydrolyzes an adenylate (AMP) moiety from 5’-end of ligation failure products and allows further attempts at completing repair (33). In our previous studies, we reported a role of Flap Endonuclease 1 (FEN1) for the processing of blocked repair intermediates with 5’-AMP (34–36). This compensatory mechanism is important to complement a deficiency in the APTX activity, which is associated with the mutations in the *aptx* gene and linked to the autosomal recessive neurodegenerative disorder Ataxia with oculomotor apraxia type 1 (AOA1) (37). Recently, the mechanism by which APTX removes 5’-AMP during the ligation of oxidative DNA damage-containing DNA ends has been suggested as a surveillance mechanism that protects LIG1 from ligation failure (32). Yet, the specificity of APTX and FEN1 for the repair intermediates including 5’-AMP and 3’-damaged or mismatched ends that mimic the ligation failure products after pol β-mediated mutagenic 8-oxodGTP or mismatch insertions is still unknown.

In this present study, we examined the importance of LIG1 fidelity for faithful substrate-product channeling and ligation of repair intermediates at the final steps of the BER pathway. For this purpose, we evaluated the LIG1 mutants with lower fidelity (E346A, E592A, and E346A/E592A) for the ligation of pol β nucleotide insertion products and the nicked DNA substrates with 3’-preinserted damaged or mismatched bases *in vitro*. Our findings revealed the mutagenic ligation of pol β 8-oxodGTP insertion products and an inefficient product channeling after pol β Watson-Crick like dGTP:T mismatch insertion by the LIG1 EE/AA mutant in a BER reaction. Moreover, the ligation efficiency of the nicked repair intermediates with preinserted 3’-8-oxodG or mismatches is dependent on the type of template base and requires the presence of double mutation at the E346 and E592 residues that reinforce high fidelity. Finally, our findings demonstrated the compensatory roles of DNA-end trimming enzymes, APTX and FEN1, for the processing of the ligation failure products with 5’-AMP in the presence of 8-oxodG or all 12 possible mismatched bases at the 3’-end of mutagenic repair intermediates. These findings herein contribute to our understanding of the efficiency and fidelity of substrate-product channeling during the final steps of BER in situations involving aberrant LIG1 fidelity and provide a novel insight into the importance of an interplay between key repair proteins for faithful BER.

## MATERIALS AND METHODS

### Preparation of DNA substrates

Oligodeoxyribonucleotides with and without a 6-carboxyfluorescein (FAM) label and the 5’-adenylate (AMP) were obtained from Integrated DNA Technologies. The DNA substrates used in this study were prepared as described previously (12–16, 34–36, 38, 39). The one nucleotide gapped DNA substrates were used for coupled assays (Supplementary Table 1). The nicked DNA substrates containing template base A, T, G, or C and 3’-preinserted bases (dA, dT, dG, or dC) or 3’-8-oxodG were used for DNA ligation assays (Supplementary Table 2). The nicked DNA substrates containing template base A, T, G, or C as well as 5’-AMP and 3’-preinserted bases (dA, dT, dG, or dC) or 3’-8-oxodG were used for APTX and FEN1 activity assays (Supplementary Table 3).

### Protein purifications

The constructs for DNA ligase I (LIG1) mutants (E346A, E592A and E346A/E592A) were prepared using the wild-type full-length DNA ligase I (pET-24b) and site-directed mutagenesis with synthetic primers and confirmed by sequencing of the coding region. The His-tag recombinant for LIG1 low fidelity mutants were purified as described previously for wild-type LIG1 with slight modifications (12–16). Briefly, the protein was overexpressed in Rosetta (DE3) pLysS *E.coli* cells (Millipore Sigma) and grown in Terrific Broth (TB) media with kanamycin (50 μgml^-1^) and chloramphenicol (34 μgml^-1^) at 37 °C. Once the OD was reached to 1.0, the cells were induced with 0.5 mM IPTG and the overexpression was continued for overnight at 16 °C. After the centrifugation, the cell was lysed in the lysis buffer (50 mM Tris-HCl pH 7.0, 500 mM NaCl, 20 mM imidazole, 10% glycerol, 1 mM PMSF, an EDTA-free protease inhibitor cocktail tablet) by sonication at 4 °C. The lysate was pelleted at 16,000xrpm for 1h at 4 °C. The cell lysis solution was filter-clarified and then loaded onto a HisTrap HP column (GE Health Sciences) that was previously equilibrated with the binding buffer A (50 mM Tris-HCl pH 7.0, 500 mM NaCl, 20 mM imidazole, 10% glycerol). The column was washed with the binding buffer A and then followed by buffer B (50 mM Tris-HCl pH 7.0, 500 mM NaCl, 35 mM imidazole, 10% glycerol). The protein was eluted with an increasing imidazole gradient (0-500 mM) of the elution buffer A at 4 °C. The collected fractions were then subsequently loaded onto HiTrap Heparin (GE Health Sciences) column that was equilibrated with binding buffer C (50 mM Tris-HCl pH 7.0, 50 mM NaCl, and 10% glycerol), and protein is eluted with elution buffer D (20 mM Tris-HCl pH 7.0, 1 M NaCl and 10% glycerol). The LIG1 protein was further purified by Resource Q and finally by Superdex 200 10/300 (GE Health Sciences) columns in the buffer (20 mM Tris-HCl pH 7.0, 200 mM NaCl, 2mM β-mercaptoethanol, and 5% glycerol).

Recombinant (GST-tagged pGEX4T3) wild-type full-length human DNA polymerase β was purified as previously described (12–16). Briefly, the protein was overexpressed in One Shot BL21(DE3)pLysS *E. coli* cells (Invitrogen) and grown at 37 °C with 0.5 mM IPTG induction. The cells were then grown overnight at 16 °C. After centrifugation, the cells were lysed at 4 °C by sonication in lysis buffer containing 1X PBS (pH 7.3) and 200 mM NaCl, and a protease inhibitor cocktail. The lysate was pelleted at 16,000 rpm for 1 h and then clarified by filtration. The pol β supernatant was loaded onto a GSTrap HP column (GE Health Sciences) that was equilibrated with 1X PBS (pH 7.3) and purified with elution buffer containing 50 mM Tris-HCl (pH 8.0), 10 mM reduced glutathione, and 1 mM DTT at 4 °C. The collected fractions were subsequently passed through a HiTrap Desalting HP column in a buffer containing 150 mM NaCl and 20 mM NaH_2_PO_4_ (pH 7.0), and then further purified by Superdex 200 Increase 10/300 chromatography (GE Healthcare). All proteins purified in this study were dialyzed against storage buffer (25 mM TrisHCl, pH 7.4, 100 mM KCl, 1 mM TCEP, and 10% glycerol), concentrated, frozen in liquid nitrogen, and stored at −80 °C. Protein quality was evaluated onto 10% SDS-PAGE, and the protein concentration was measured using absorbance at 280 nm.

### Coupled assays

The coupled assays were performed to measure pol β and LIG1 activities simultaneously in the same reaction mixture as described previously (12–16, 38, 39). For this purpose, we used the one nucleotide gap DNA substrates (Supplementary Table 1). Briefly, the reaction was initiated by the addition of pol β (10 nM) and LIG1 (10 nM) to a mixture containing 50 mM Tris-HCl (pH 7.5), 100 mM KCl, 10 mM MgCl2, 1 mM ATP, 1 mM DTT, 100 μg ml^-1^ BSA, 10% glycerol, the DNA substrate (500 nM), and 8-oxodGTP or dGTP (100 μM). The reaction mixture was incubated at 37 °C and stopped at the indicated time points in the figure legends. The reaction products were quenched with addition of gel loading buffer (95% formamide, 20 mM ethylenediaminetetraacetic acid, 0.02% bromophenol blue and 0.02% xylene cyanol) and then separated by electrophoresis on an 18% polyacrylamide gel. The gels were scanned with a Typhoon PhosphorImager (Amersham Typhoon RGB), and the data were analyzed using ImageQuant software. The coupled assays were performed similarly for wild-type and low fidelity LIG1 mutants E346A, E592A or E346A/E592A.

### DNA ligation assays

The ligation assays were performed to analyze the substrate specificity of LIG1 as described previously (12–16, 38, 39). For this purpose, we used the nicked DNA substrates with 3’-preinserted 8-oxodG or mismatched bases (Supplementary Table 2). Briefly, the reaction was performed in a mixture containing 50 mM Tris-HCl (pH 7.5), 100 mM KCl, 10 mM MgCl2, 1 mM ATP, 1 mM DTT, 100 μg ml^-1^ BSA, 10% glycerol, the nicked DNA substrate (500 nM), and LIG1 (100 nM). The reaction mixtures were then incubated at 37 °C for the times indicated in the figure legends and were quenched by mixing with an equal volume of loading dye. The products were separated, and the data were analyzed as described above. The ligation assays were performed similarly for wild-type and low fidelity LIG1 mutants E346A, E592A or E346A/E592A.

### Aprataxin and FEN1 assays

Aprataxin (APTX) and flap endonuclease 1 (FEN1) activity analyses were performed as described previously (34–36). For this purpose, we used the nicked DNA substrates containing 5’-AMP and 3’-preinserted damaged (8-oxodG) or all possible 12 mismatched bases (Supplementary Table 3). Briefly, APTX assay was performed in a mixture containing 50 mM Tris-HCl (pH 7.5), 40 mM KCl, 5 mM EDTA, 1 mM DTT, 5% glycerol, and the nicked DNA substrate (500 nM). FEN1 assay was performed in a mixture containing 50 mM Hepes (pH 7.5), 20 mM KCl, 0.5 mM EDTA, 2 mM DTT, 10 mM MgCl_2_, and the nicked DNA substrate (500 nM). For both assays, the reactions were initiated with the addition of APTX (100 nM) or FEN1 (100-500 nM) and incubated at 37 °C. The activity assays were stopped at 15 min for FEN1 and at the indicated time points in the figure legends for APTX by mixing with an equal volume of loading dye. The reaction products were then analyzed as described above.

### Structure modelling

Structure analysis was performed based on the crystal structure of LIG1 (PDB:6P0E) using the Coot software (40). All structural images were drawn using PyMOL (http://www.pymol.org/).

## RESULTS

### Low fidelity DNA ligase I stimulates the mutagenic ligation of pol β 8-oxodGTP insertion products

We previously demonstrated that the nicked repair product of pol β 8-oxodGTP insertion cannot be used as a substrate by DNA ligase I (LIG1) during substrate-product channeling at the downstream steps of BER pathway (12–16). In the present study, we first evaluated the effect of low fidelity LIG1 for the ligation of pol β 8-oxodGTP insertion products *in vitro*. For this purpose, we used the coupled assay that measures the activities of pol β and LIG1 simultaneously in the reaction mixture including LIG1 wild-type or LIG1 E346A/E592A (EE/AA) mutant, pol β, 8-oxodGTP, and the one nucleotide gap DNA substrate with template base A or C (Figure 1A).

**Figure 1.**
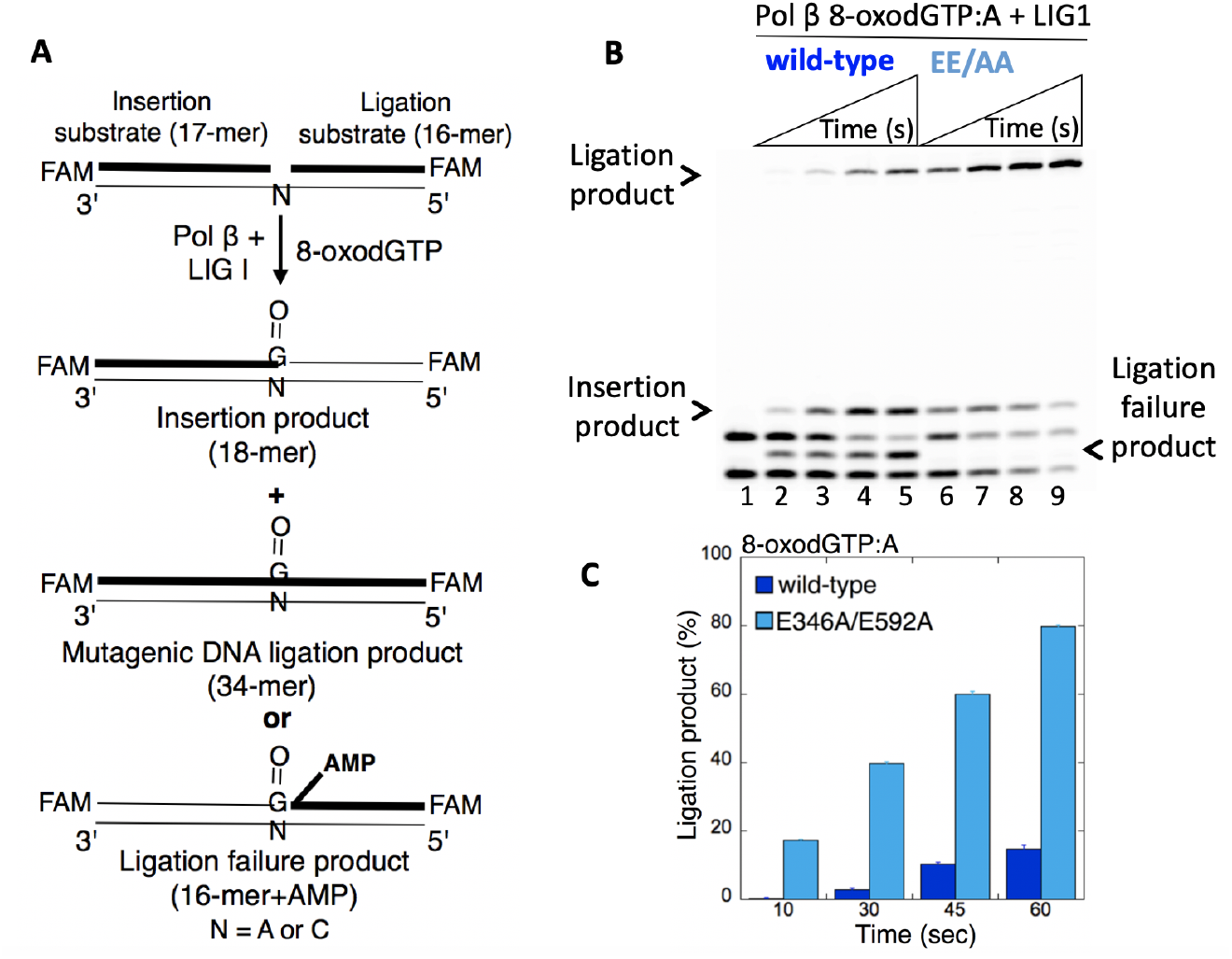
Mutagenic ligation of pol β 8-oxodGTP insertion opposite A by low fidelity LIGI. (**A**) Illustrations of the one nucleotide gapped DNA substrate and the products of insertion, ligation, and ligation failure obtained in the coupled reaction. (**B**) Lane 1 is the negative enzyme control of the one nucleotide gapped DNA substrate with template base A. Lanes 2-5 and 6-9 show the insertion coupled to ligation products by LIGI wild type and E346A/E592A mutant, respectively, obtained at the time points 10, 30, 45, and 60 sec. (**C**) The graph shows the time-dependent changes in the ligation products and the data are presented as the averages from three independent experiments ± SDs.

In consistent with our previous studies, we observed that wild-type LIG1 cannot ligate the products of pol β 8-oxodGTP insertion opposite A (Figure 1B, lanes 2-5). This results in the ligation failure and accumulation of abortive ligation products with 5’-adenylate (5’-AMP). On the other hand, there was no ligation failure after pol β oxidized nucleotide insertions in the presence of the LIG1 mutant EE/AA that interferes with ligase fidelity. In this case, we observed the mutagenic ligation of pol β 8-oxodGTP:A insertion product (Figure 1B, lanes 6-9). The amount of this ligation product was ~10-fold higher when compared to those obtained with the wild-type LIG1 (Figure 1C). Similarly, pol β 8-oxodGTP:C insertion products can be efficiently ligated in the coupled reaction including the LIG1 mutant EE/AA (Figure 2A, lanes 9-15) in contrast to the ligation failure with the wild-type LIG1 (Figure 2A, lanes 2-8). The amount of this mutagenic ligation products was higher than those of the wild-type LIG1 (Figure 2B), but lower than the ligation after pol β 8-oxodGTP:A insertion by the LIG1 mutant EE/AA (Supplementary Figure 1).

**Figure 2.**
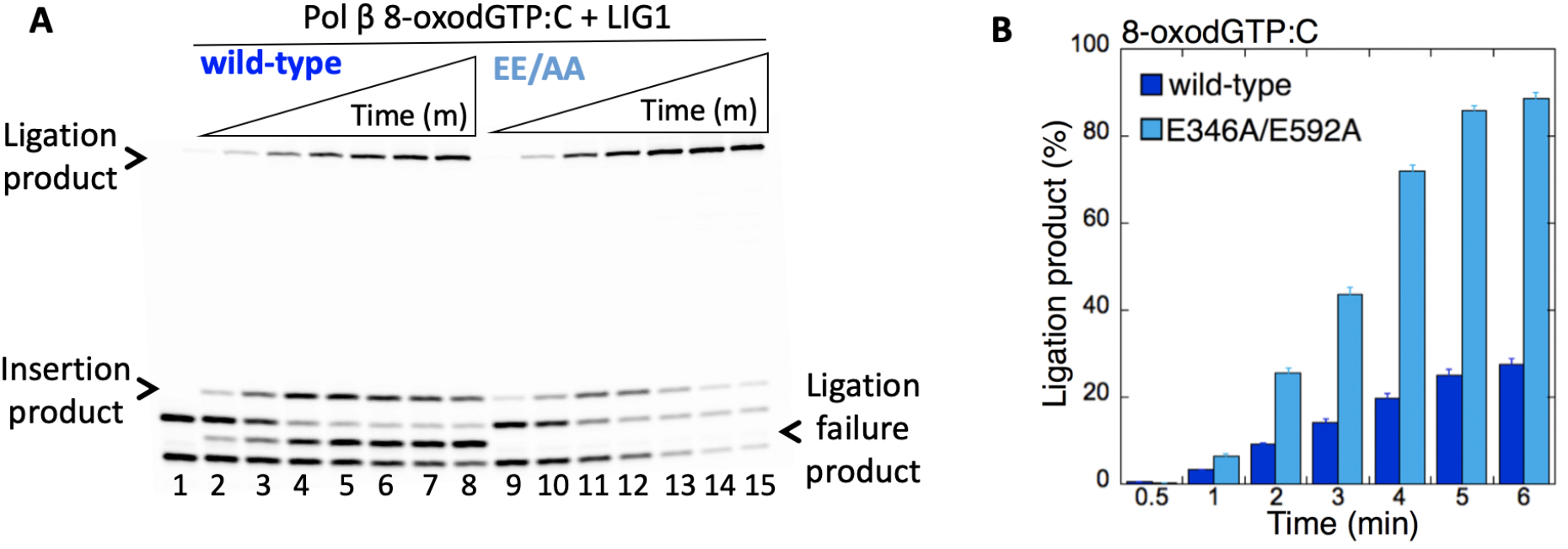
Mutagenic ligation of pol β 8-oxodGTP insertion opposite C by low fidelity LIG1. (**A**) Lane 1 is the negative enzyme control of the one nucleotide gapped DNA substrate with template base C. Lanes 2-8 and 9-15 show the insertion coupled to ligation products by LIGI wild type and E346A/E592A mutant, respectively, obtained at the time points 0.5, 1, 2, 3, 4, 5, and 6 min. (**B**) The graph shows the time-dependent changes in the ligation products and the data are presented as the averages from three independent experiments ± SDs.

In the control coupled reactions including pol β, LIG1, dGTP, and one nucleotide gap DNA substrate with template base C, we also evaluated the ligation of pol β dGTP:C insertion products in the presence of wild-type vs the LIG1 mutant EE/AA (Figure 3A). The results demonstrated the conversion of pol β correct nucleotide insertions to complete ligation products over the time of reaction incubation for both the wild-type and low fidelity LIG1 mutant (Figure 3B, lanes 2-8 and 9-15, respectively). In this case, we did not observe significant difference in the amount of ligation products between wild-type and the LIG1 mutant EE/AA (Figure 3C), while the substrate-product channeling from pol β to LIG1 was faster (*i.e*., a decrease in the amount of the dGTP:C insertion products, Figure 3D) in the presence of the low fidelity mutant. This could be due to the facilitated hand off of repair intermediate with a correct base inserted by pol β to the next DNA ligation step of the repair pathway.

**Figure 3.**
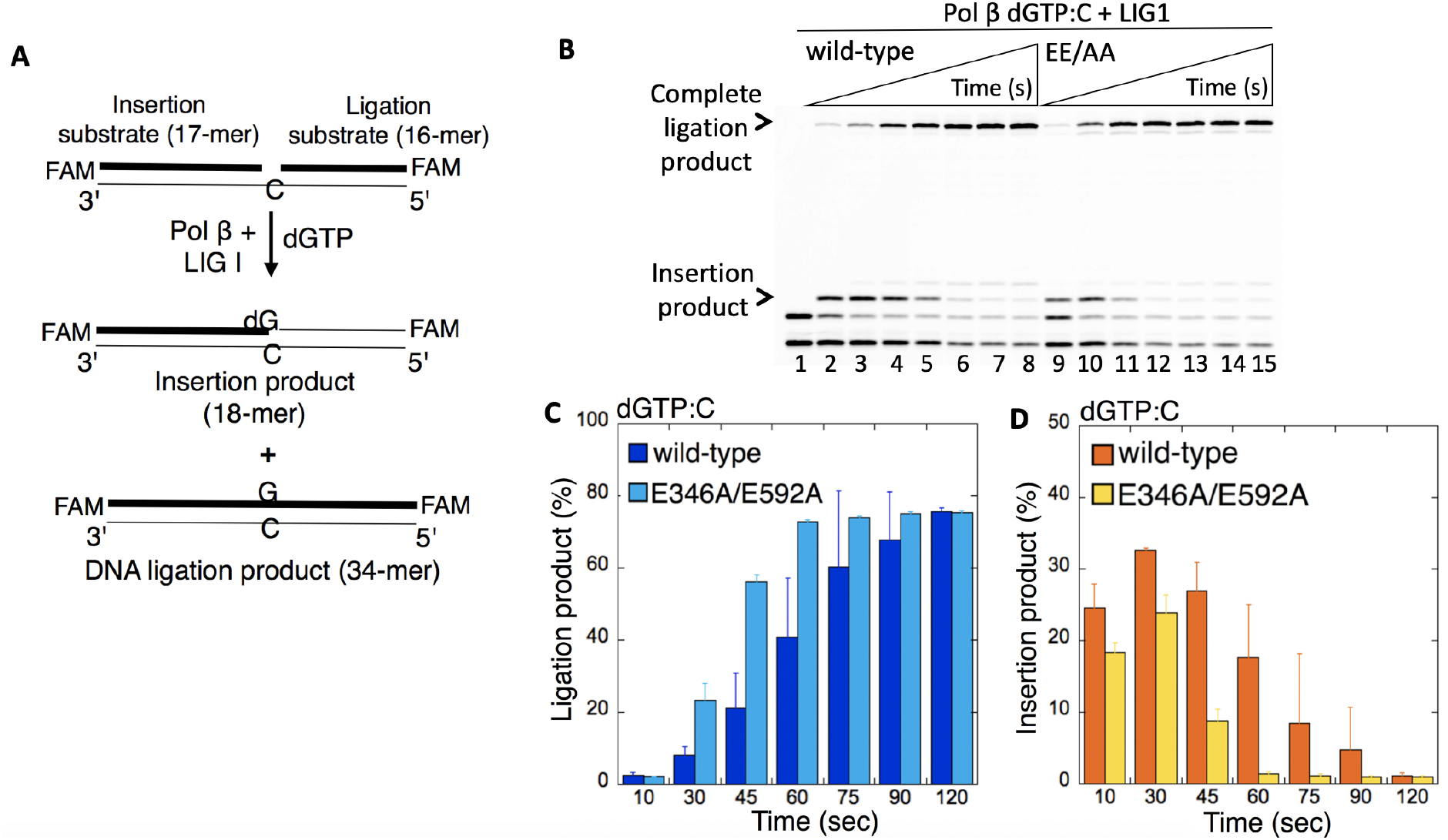
Ligation of pol β dGTP insertion opposite C by low fidelity LIGI. (**A**) Illustrations of the one nucleotide gapped DNA substrate and the products of insertion and ligation obtained in the control coupled reaction. (**B**) Lane 1 is the negative enzyme control of the one nucleotide gapped DNA substrate with template base C. Lanes 2-8 and 9-15 show the insertion coupled to ligation products by LIGI wild type and E346A/E592A mutant, respectively, obtained at the time points 10, 30, 45, 60, 75, 90, and 120 sec. (**C,D**) The graphs show the time-dependent changes in the products of insertion and ligation. The data are presented as the averages from three independent experiments ± SDs.

### Mutagenic ligation of the nicked repair intermediates with preinserted 3’-8-oxodG by low fidelity DNA ligase I

We then evaluated the importance of the ligase fidelity on DNA end-joining of the repair intermediates that mimic DNA polymerase oxidized nucleotide (8-oxodGTP) insertion products *in vitro*. For this purpose, we used the ligation assay in a reaction mixture including LIG1 wild-type or low fidelity mutants (E346A, E592A, or E346A/E592A) and the nicked DNA substrates including preinserted 3’-8oxodG opposite template base A, T, G, or C (Figure 4B).

**Figure 4.**
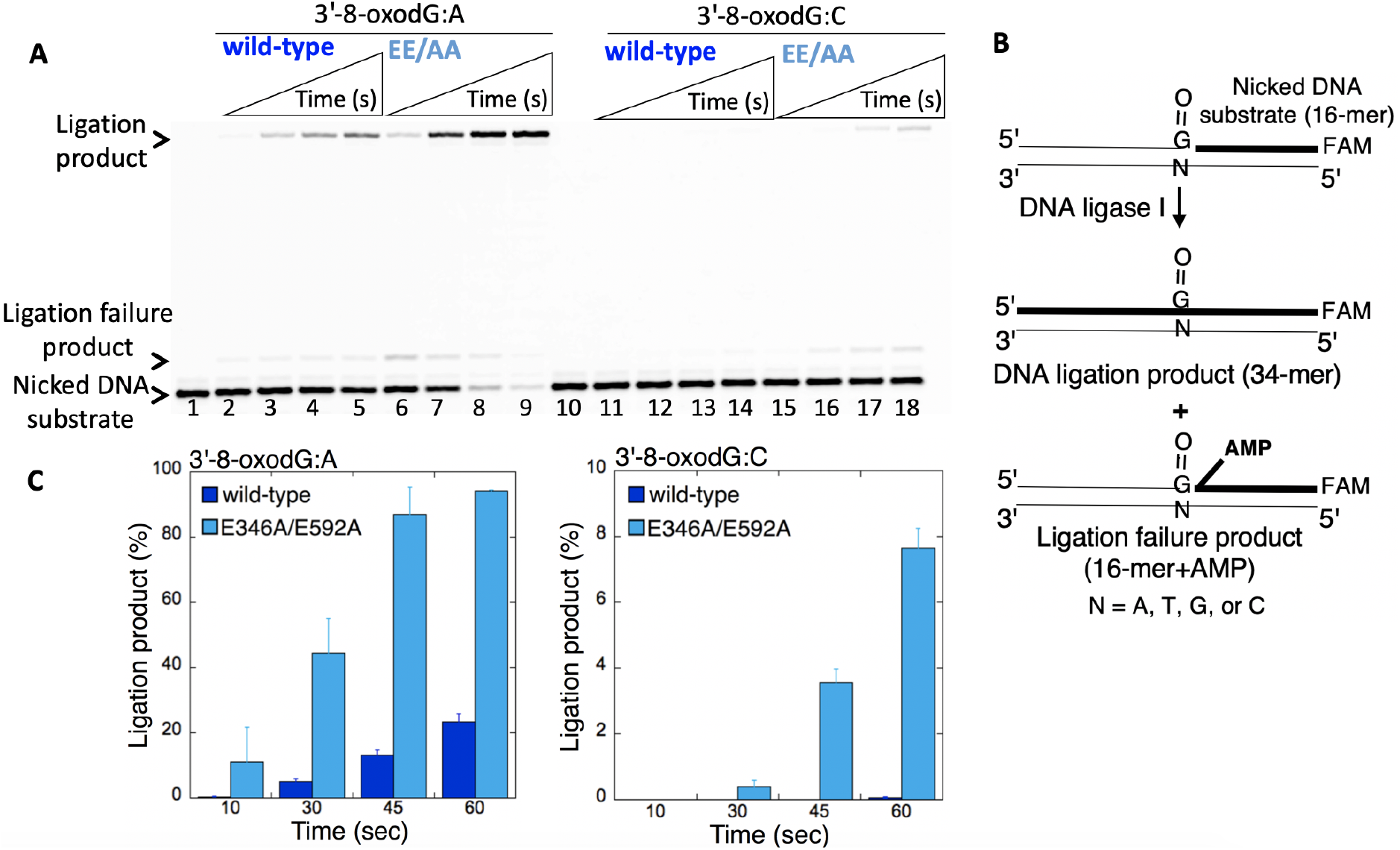
Ligation efficiency of the repair intermediate with preinserted 3’-8-oxodG by low fidelity LIG1. (**A**) Lanes 1 and 10 are the negative enzyme controls of the nicked DNA substrates with 3’-8-oxodG opposite template base A or C, respectively. Lanes 2-5 and 6-9 show the ligation products by LIGI wild type and E346A/E592A mutant, respectively, for 3’-8-oxodG:A substrate, obtained at the time points 10, 30, 45, and 60 sec. Lanes 11-14 and 15-18 show the ligation products by LIGI wild type and E346A/E592A mutant, respectively, for 3’-8-oxodG:C substrate, obtained at the time points 10, 30, 45, and 60 sec. (**B**) Illustrations of the nicked DNA substrate with 3’-8-oxodG and the products observed in the ligation reaction. (**C**) The graphs show the time-dependent changes in the ligation products and the data are presented as the averages from three independent experiments ± SDs.

The results showed the mutagenic ligation of the nicked DNA substrates with 3’-8-oxodG:A (Figure 4A, lanes 6-9) and 3’-8-oxodG:C (Figure 4A, lanes 15-18) by the LIG1 mutant EE/AA in comparison to an inefficient ligation by wild-type enzyme (Figure 4A, lanes 2-5 and 11-14, respectively). For both DNA substrates, the mutagenic ligation was enhanced by the presence of the EE/AA mutation (Figure 4C).

We then tested the contribution of each of ligase active site residues (E346 and E592) to the ligation efficiency of repair intermediates that mimic pol β oxidized nucleotide insertion products using the ligation reactions at longer time points (0.5-10 min). For the nicked DNA substrate with preinserted 3’-8-oxodG opposite A, we did not observe a significant difference between LIG1 single mutants E346A (Figure 5A, lanes 2-7) and E592A (Figure 5A, lanes 8-13) vs the LIG1 double mutant EE/AA (Figure 5A, lanes 14-19). The amount of mutagenic ligation products was similar (Figure 5B). For the nicked DNA substrate with preinserted 3’-8-oxodG opposite C, we obtained the mutagenic ligation products that were enhanced by the presence of the LIG1 double mutant EE/AA (Figure 5C, lanes 14-19) when compared to the ligation by E346A (Figure 5C, lanes 2-7) and E592A (Figure 5C, lanes 8-13). There was ~80-fold difference in the amount of ligation products between wild-type and the EE/AA mutant (Figure 5D).

**Figure 5.**
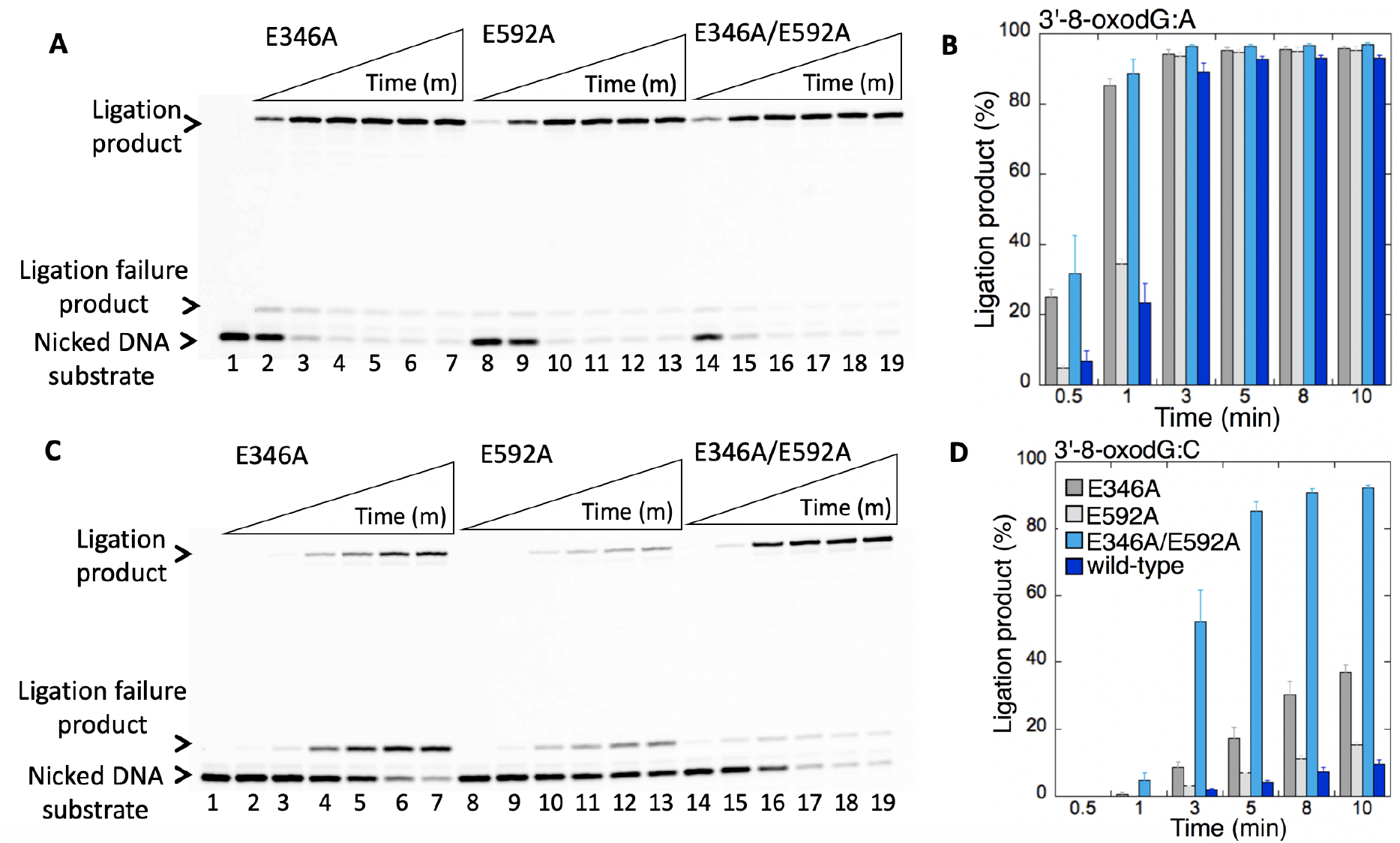
Ligation efficiency of the repair intermediate with 3’-8-oxodG opposite A or C by low fidelity LIG1. (**A,C**) Lane 1 is the negative enzyme control of the nicked DNA substrate with 3’-8-oxodG opposite template base A (A) or C (C). Lanes 2-7, 8-13, and 14-19 show the ligation products by LIGI mutants E346A, E592A, and E346A/E592A, respectively, for 3’-8-oxodG:A (A) and 3’-8-oxodG:C (C) substrates, obtained at the time points 0.5, 1, 3, 5, 8, and 10 min. (**B,D**) The graphs show the time-dependent changes in the ligation products and the data are presented as the averages from three independent experiments ± SDs.

Interestingly, for the nicked DNA substrates containing 3’-8-oxodG opposite G or T, we did not observe mutagenic DNA ligation in the presence of any LIG1 mutant with perturbed fidelity (Figure 6). There was only ligation failure products on nicked DNA with 3’-8-oxodG:G ends by LIG1 E346A (Figure 6A, lanes 2-7), E592A (Figure 6A, lanes 8-13), and E346A/E592A (Figure 6A, lanes 14-19). Similarly, for the nicked DNA substrate with 3’-8-oxodG opposite T, we obtained a negligible amount of ligation products only by LIG1 E346A/E592A (Figure 6D). On the other hand, the results showed accumulation of ligation failure products in the presence of E346A (Figure 6C, lanes 2-7), E592A (Figure 6C, lanes 8-13) and E346A/E592A (Figure 6C, lanes 14-19) as revealed by the formation of the 5’-adenylate product, *i.e*., addition of AMP to the 5’-end of the substrate. When compared with the ligation of the preinserted 3’-8-oxodG for all four possible template bases by wild-type LIG1 (Supplementary Figure 2), overall results indicate that the mutagenic ligation of the nicked repair intermediates with preinserted 3’-8-oxodG by low fidelity LIG1 is template base-dependent and requires the mutations at both ligase active site residues (E346 and E592) that reinforce the ligase fidelity (Supplementary Figure 3).

**Figure 6.**
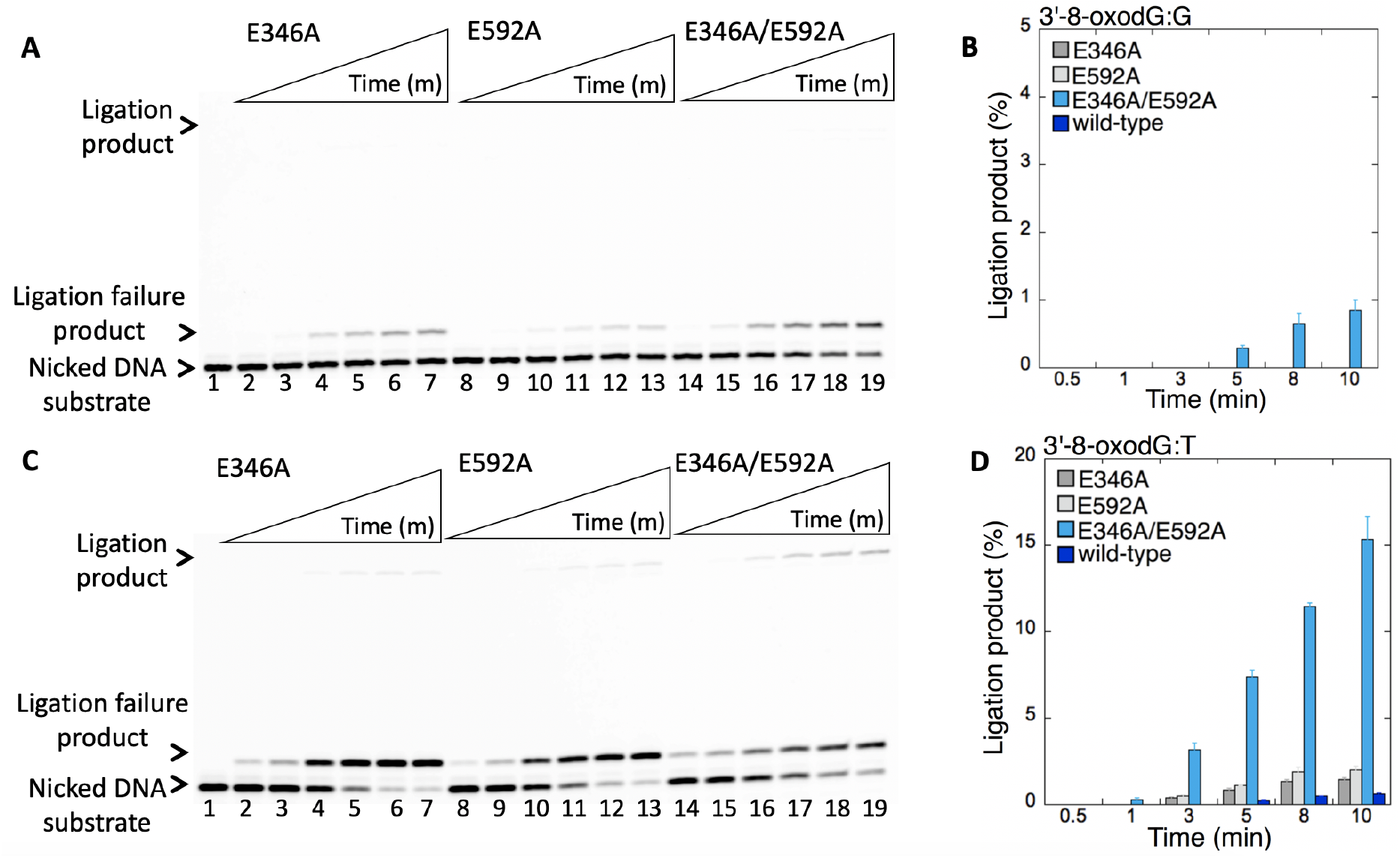
Ligation efficiency of the repair intermediate with 3’-8-oxodG opposite G or T by low fidelity LIG1. (**A,C**) Lane 1 is the negative enzyme control of the nicked DNA substrate with 3’-8-oxodG opposite template base G (A) or T (C). Lanes 2-7, 8-13, and 14-19 show the ligation products by LIGI mutants E346A, E592A, and E346A/E592A, respectively, for 3’-8-oxodG:G (A) and 3’-8-oxodG:T (C) substrates, obtained at the time points 0.5, 1, 3, 5, 8, and 10 min. (**B,D**) The graphs show the time-dependent changes in the ligation products and the data are presented as the averages from three independent experiments ± SDs.

### Inefficient substrate-product channeling from pol β Watson-Crick like dG:T insertion to low fidelity DNA ligase I

In our prior work, we reported that LIG1 efficiently ligates the pol β Watson-Crick like dGTP:T insertion products (16). In order to further examine the effect of staggered LIG1 conformation with perturbed fidelity on the substrate-product channeling of repair intermediates at the downstream steps of the BER pathway, we also evaluated the ligation of pol β dGTP:T insertion products in the coupled reaction including LIG1 wild-type or E346A/E592A mutant (EE/AA), pol β, dGTP, and the one nucleotide gap DNA substrate with template base T (Figure 7A).

**Figure 7.**
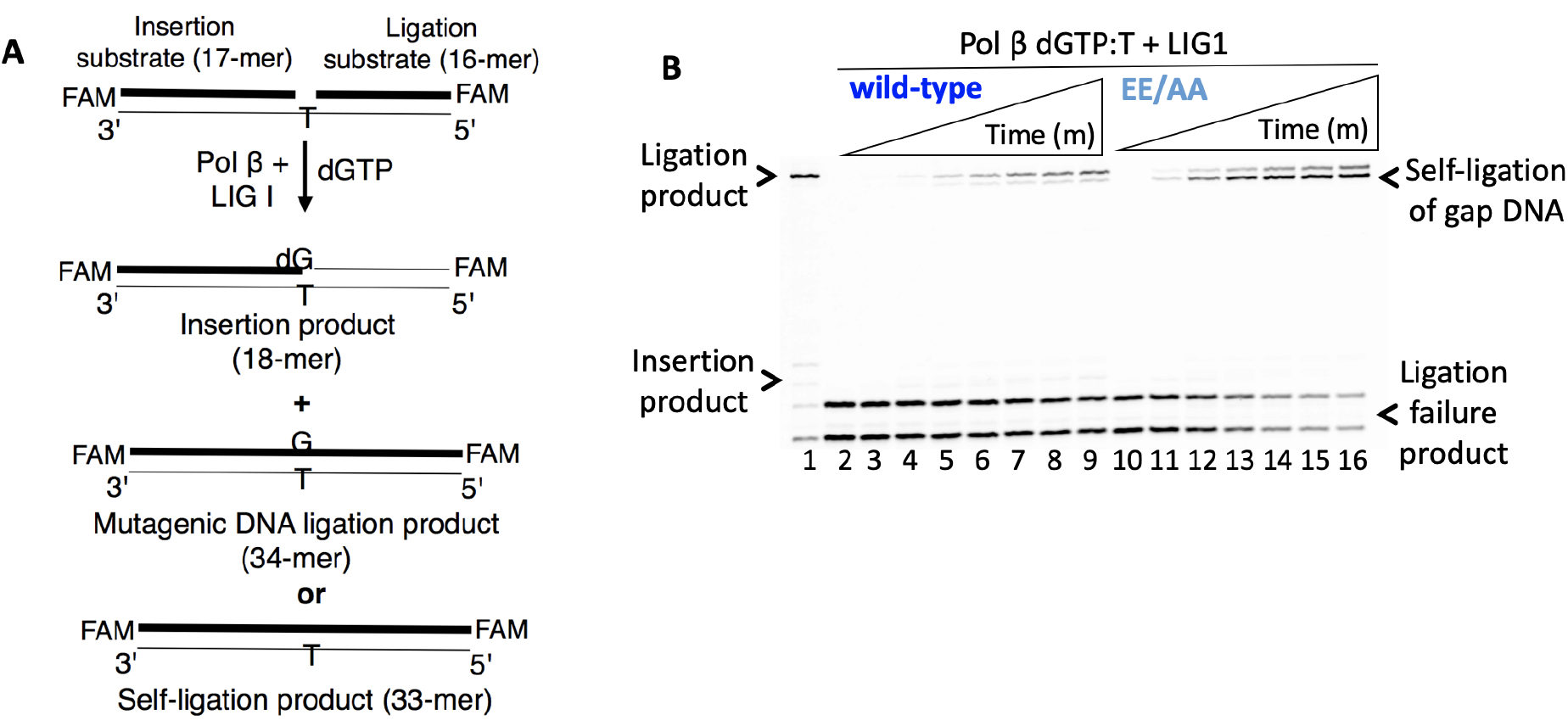
Ligation of pol β dGTP insertion opposite T by low fidelity LIGI. (**A**) Illustrations of the one nucleotide gapped DNA substrate and the products obtained in the control coupled reaction. (**B**) Lane 1 is the positive control for the ligation of pol β dGTP:C insertion product by wild-type LIG1. Lane 2 is the negative enzyme control of the one nucleotide gapped DNA substrate with template base T. Lanes 3-9 and 10-16 show the insertion coupled to ligation products by LIGI wild type and E346A/E592A mutant, respectively, obtained at the time points 10, 30, 45, 60, 75, 90, and 120 sec. The gel is a representative of three independent experiments.

Surprisingly, the products of mismatch insertion coupled to ligation by the LIG1 mutant EE/AA were mainly the self-ligation products of the one nucleotide gapped DNA itself (Figure 7B, lanes 10-16), as revealed by the difference in the size of the products with the ligation of the pol β dG:T mismatch insertion (Figure 7B, lanes 3-9 vs 10-16) and the ligation of pol β dG:C correct insertion (Figure 7B, lane 1 vs 10-16) by the wild-type LIG1. We then tested the ligation of nicked DNA substrate with preinserted 3’-dG:T mismatch in a ligation reaction (Figure 8B). In contrast to the coupled reaction including pol β and LIG1, the results revealed mutagenic ligation of the nicked DNA substrate with 3’-dG:T mismatch, which is stimulated by low fidelity LIG1 mutants E346A (Figure 8B, lanes 6-9), E592A (Figure 8B, lanes 10-13), and E346A/E592A (Figure 8B, lanes 14-17). There was ~ 50-fold increase in the amount of ligation products between the wild-type LIG1 and E346A/E592A (Figure 8C). The difference in the ligation efficiency we observed between the coupled (after pol β mismatch insertion into a gap) vs ligation (end joining of nicked DNA by LIG1 alone) reactions could be due to the interplay between these two key repair enzymes at the downstream steps of the BER pathway, which may affect the repair outcomes.

**Figure 8.**
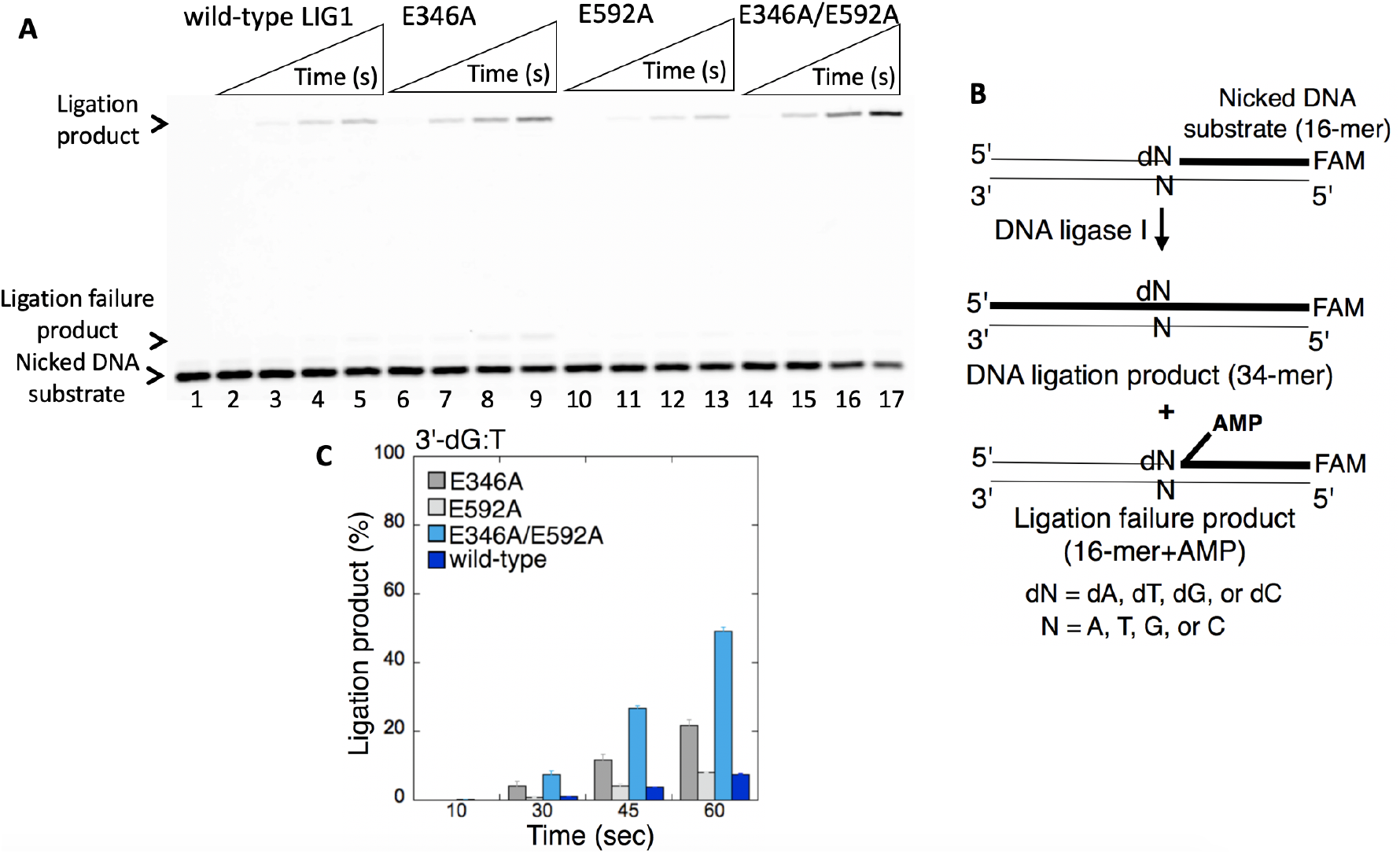
Ligation efficiency of the repair intermediate with Watson-Crick-like 3’-dG:T mismatch by low fidelity LIG1. (**A**) Lane 1 is the negative enzyme control of the nicked DNA substrate with 3’-dG:T mismatch, and lanes 2-5, 6-9, 10-13, and 14-17 show the ligation products by LIGI wild type, E346A, E592A, and E346A/E592A mutants, respectively, obtained at the time points 10, 30, 45, and 60 sec. (**B**) Illustrations of the nicked DNA substrate and the ligation and ligation failure products obtained in the reaction including 3’-preinserted mismatches. (**C**) The graph shows the time-dependent changes in the ligation products and the data are presented as the averages from three independent experiments ± SDs.

### The effect of aberrant LIG1 fidelity on DNA end-joining of 3’-preinserted non-canonical mismatches

We previously reported that the BER DNA ligases (LIG1 and LIG3) recognize subtle base differences at either the template or the 3’-end when sealing nicked DNA, and the mismatch discrimination of the ligases can vary depending on the DNA end structure of the repair intermediate (16). In the present study, we evaluated the 3’-end surveillance of LIG1 with low fidelity for the ligation of 3’-preinserted mismatches that mimic pol β mismatch insertion products. For this purpose, we tested the nicked DNA substrates that wild-type LIG1 exhibits compromised ligation, *i.e*., 3’-preinserted dA:G, dG:A, dC:C, dT:T (Figure 8B).

For the nicked substrates with DNA ends containing purine-purine base mismatches 3’-dA:G and 3’-dG:A (Figure 9), we did not observe ligation products by LIG1single mutants E346A (Figures 9A and 9C, lanes 6-9) and E592A (Figures 9A and 9C, lanes 10-13). This was similar to the inefficient ligation by wild-type LIG1 (Figures 9A and 9C, lanes 2-5). Yet, the presence of double mutation (Figures 9A and 9C, lanes 14-17) stimulated the mutagenic ligation and there was ~ 40- to 80-fold difference between wild-type and E346A/E592A mutant (Figures 9B and 9D). For the nicked substrates with DNA ends containing pyrimidine pairs 3’-dC:C and 3’-dT:T (Figure 10), in comparison with an inefficient ligation by the wildtype LIG1 (Figures 10A and 10C, lanes 2-5), we obtained the mutagenic ligation products by LIG1 mutants E346A (Figures 10A and 10C, lanes 6-9), E592A (Figures 10A and 10C, lanes 10-13), and E346A/E592A (Figures 10A and 10C, lanes 14-17). For both DNA substrates, the presence of double mutation significantly enhanced mutagenic ligation and there was ~ 80-fold difference in the amount of ligation products between wild-type and E346A/E592A (Figures 10B and 10D). Overall results indicate that the mutagenic ligation of repair intermediates with preinserted 3’-mismatched bases by low-fidelity LIG1 is dependent of the architecture of DNA ends and requires amino acid changes at both glutamic acid E346 and E592 residues (Supplementary Figure 4). In the control ligation reactions including the nicked DNA substrate with preinserted 3’-dG:C correctly base-paired ends, we confirmed the efficient ligation by LIG1wild-type and low-fidelity mutants (Supplementary Figure 5).

**Figure 9.**
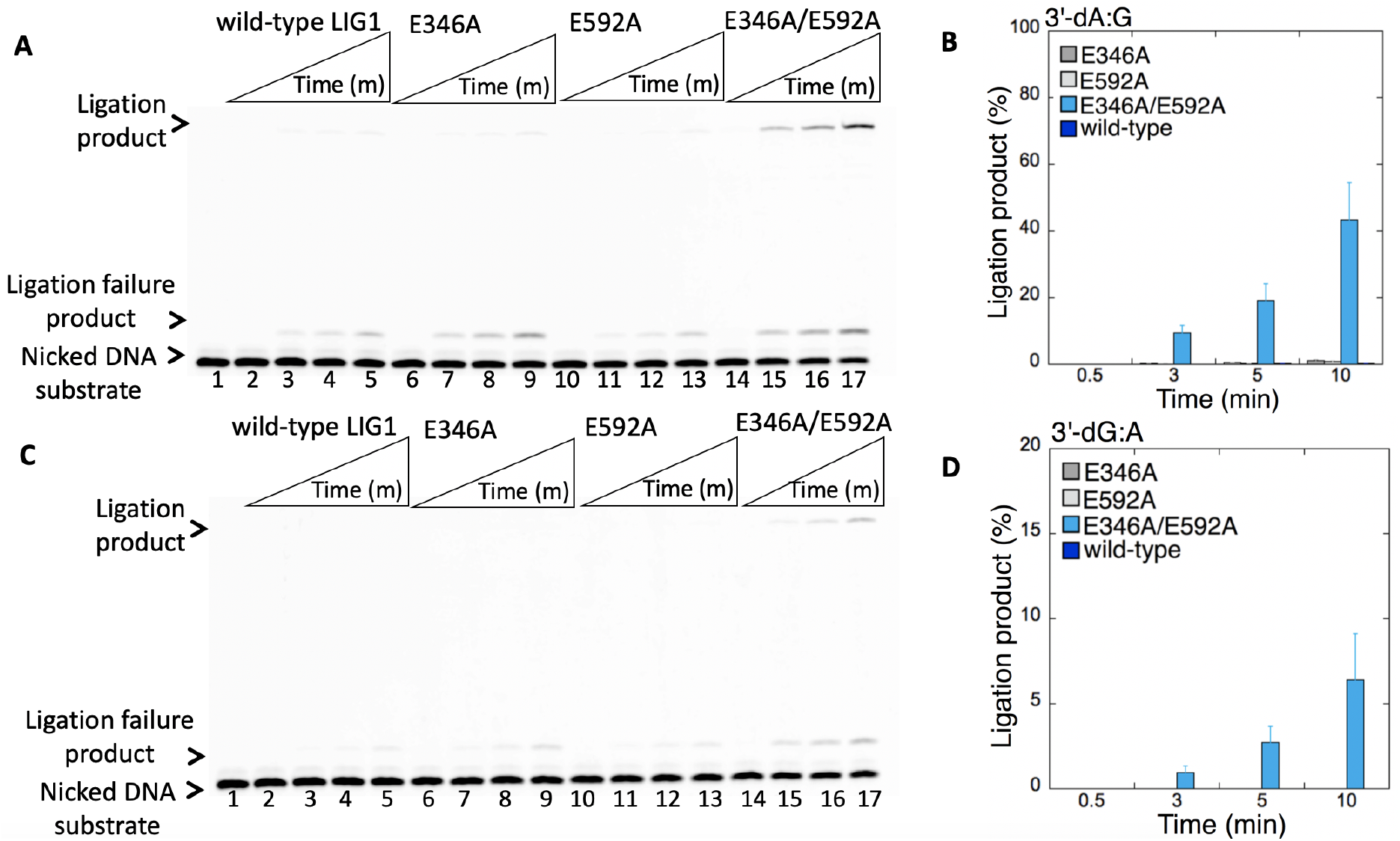
Ligation efficiency of the repair intermediates with 3’-preinserted dA:G and dG:A mismatches by low fidelity LIG1. (**A,C**) Lane 1 is the negative enzyme control of the nicked DNA substrate with 3’-dA:G (A) and 3’-dG:A (C) mismatches. Lanes 2-5, 6-9, 10-13 and 14-17 show the reaction products by LIG1 wild type, E346A, E592A, and E346A/E592A mutants, respectively, for 3’-dA:G (A) and 3’-dG:A (C) substrates, obtained at the time points 0.5, 3, 5 and 10 min. (**B**,**D)** The graphs show the time-dependent changes in the ligation products and the data are presented as the averages from three independent experiments ± SDs.

**Figure 10.**
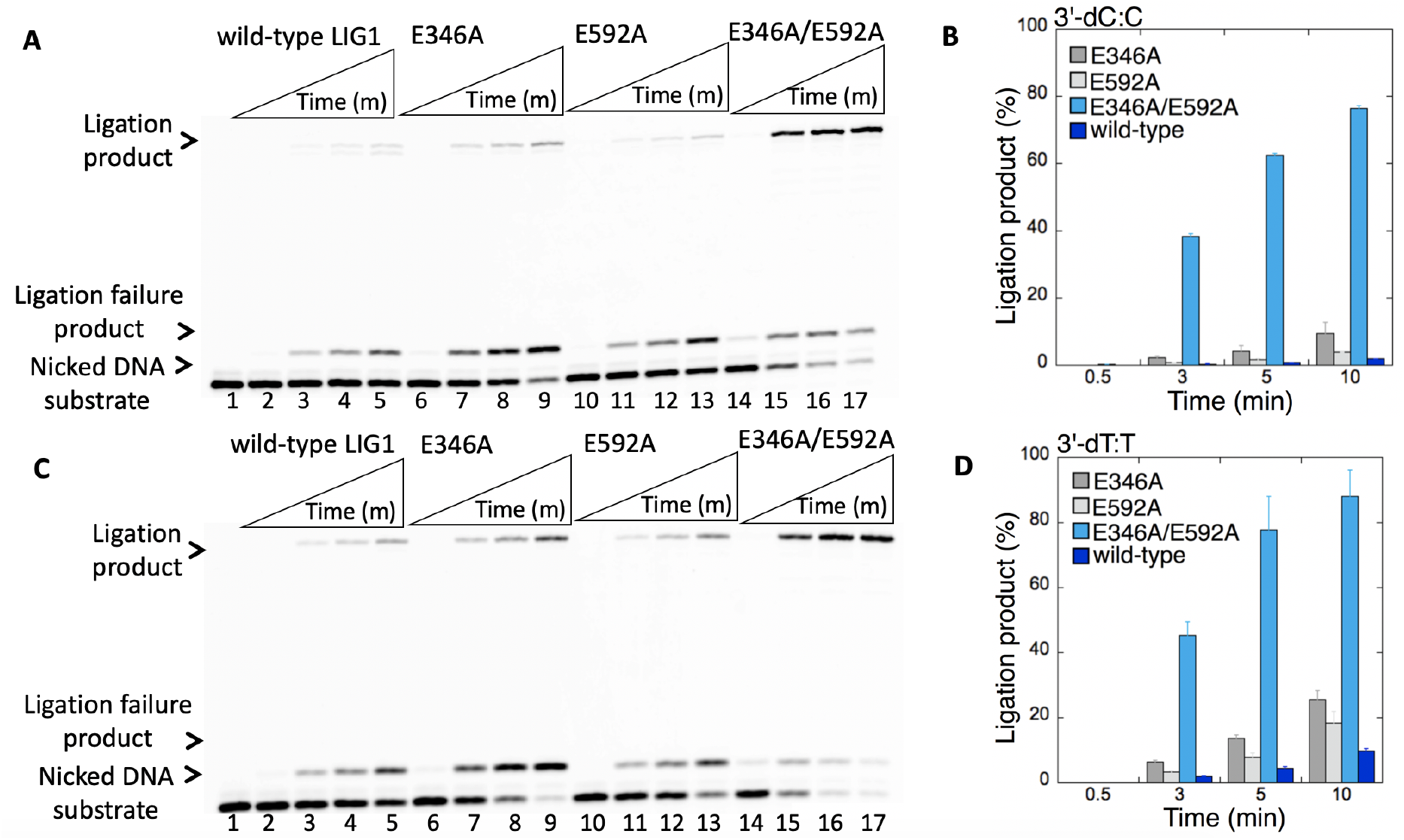
Ligation efficiency of the repair intermediates with 3’-preinserted dC:C and dT:T mismatches by low fidelity LIG1. (**A,C**) Lane 1 is the negative enzyme control of the nicked DNA substrate with 3’-dC:C (A) and 3’-dT:T (C) mismatches. Lanes 2-5, 6-9, 10-13 and 14-17 show the reaction products by LIGI wild type, E346A, E592A, and E346A/E592A mutants, respectively, for 3’-dC:C (A) and 3’-dT:T (C) substrates, obtained at the time points 0.5, 3, 5 and 10 min. (**B**,**D)** The graphs show the time-dependent changes in the ligation products and the data are presented as the averages from three independent experiments ± SDs.

### Roles of APTX and FEN1 in the processing of ligation failure intermediates with 5’-adenylate and 3’-8-oxodG or mismatch

We finally examined the DNA end-processing proteins APTX and FEN1 for their functions to clean 5’-ends from the repair intermediates containing an adenylate (AMP) block that mimic the ligation failure products after pol β 8-oxodGTP or mismatch insertions. For this purpose, we evaluated their enzymatic activities in a reaction mixture including the nicked DNA substrates with preinserted 5’-AMP and 3’-preinserted 8oxodG or all possible 12 non-canonical base pairs *in vitro*.

With the nicked DNA substrates with 5’-AMP and 3’-8-oxodG (Figures 11C and 12C), we observed an efficient removal of 5’-adenylate block by APTX from DNA ends including 3’-8-oxodG opposite template base A (Figure 11A, lanes 3-8) and template base C (Figure 11A, lanes 10-15). The results showed no significant difference in the amount of 5’-AMP removal products from both DNA substrates (Figure 11B). For FEN1 activity, we obtained the products of both 5’-AMP removal and nucleotide excisions from the nicked DNA substrates containing 3’-8-oxodG opposite template base A (Figure 12A, lanes 3-5) and template base C (Figure 12A, lanes 7-9). The amount of nucleotide excision products showed no significant difference and increased as a function of FEN1 concentration (Figure 12B).

**Figure 11.**
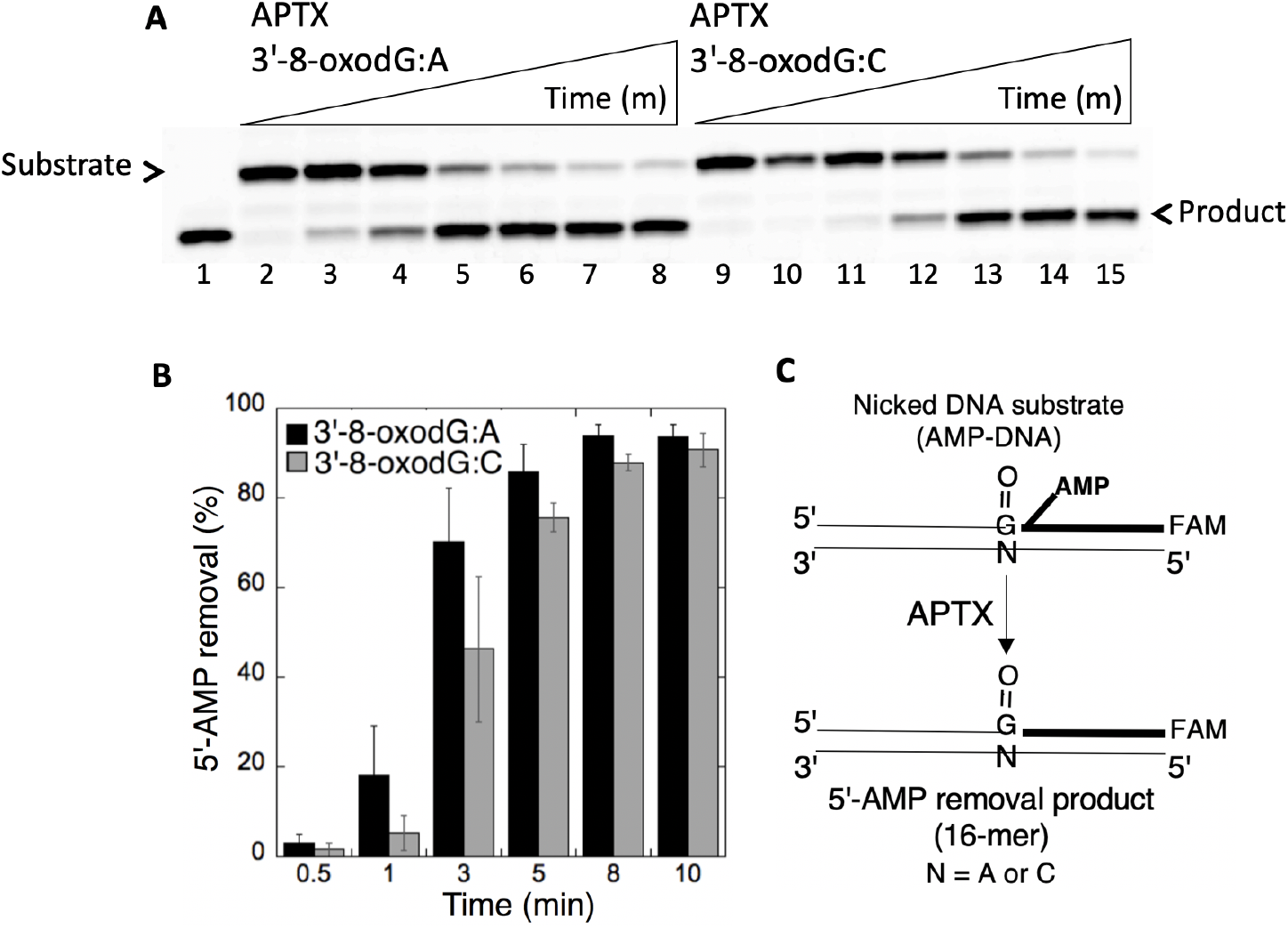
Removal of 5’-AMP by APTX from the ligation failure products with 3’-8-oxodG. (**A**) Lane 1 is a size marker that corresponds to the oligonucleotide without an AMP moiety. Lanes 2 and 9 are the negative enzyme controls of the nicked DNA substrates with 5’-AMP and 3’-8-oxodG opposite template base A or C, respectively. Lanes 3-8 and 10-15 show the products of 5’-AMP removal from 3’-8-oxodG:A and 3’-8-oxodG:C substrates, respectively, obtained at the time points 0.5, 1, 3, 5, 8, and 10 min. (**B**) The graph shows the time-dependent changes in the products of 5’-AMP removal and the data are presented as the averages from three independent experiments ± SDs. (**C**) Illustrations of the nicked DNA substrate with 5’-AMP and 3’-8-oxodG and the products observed in the APTX reaction.

**Figure 12.**
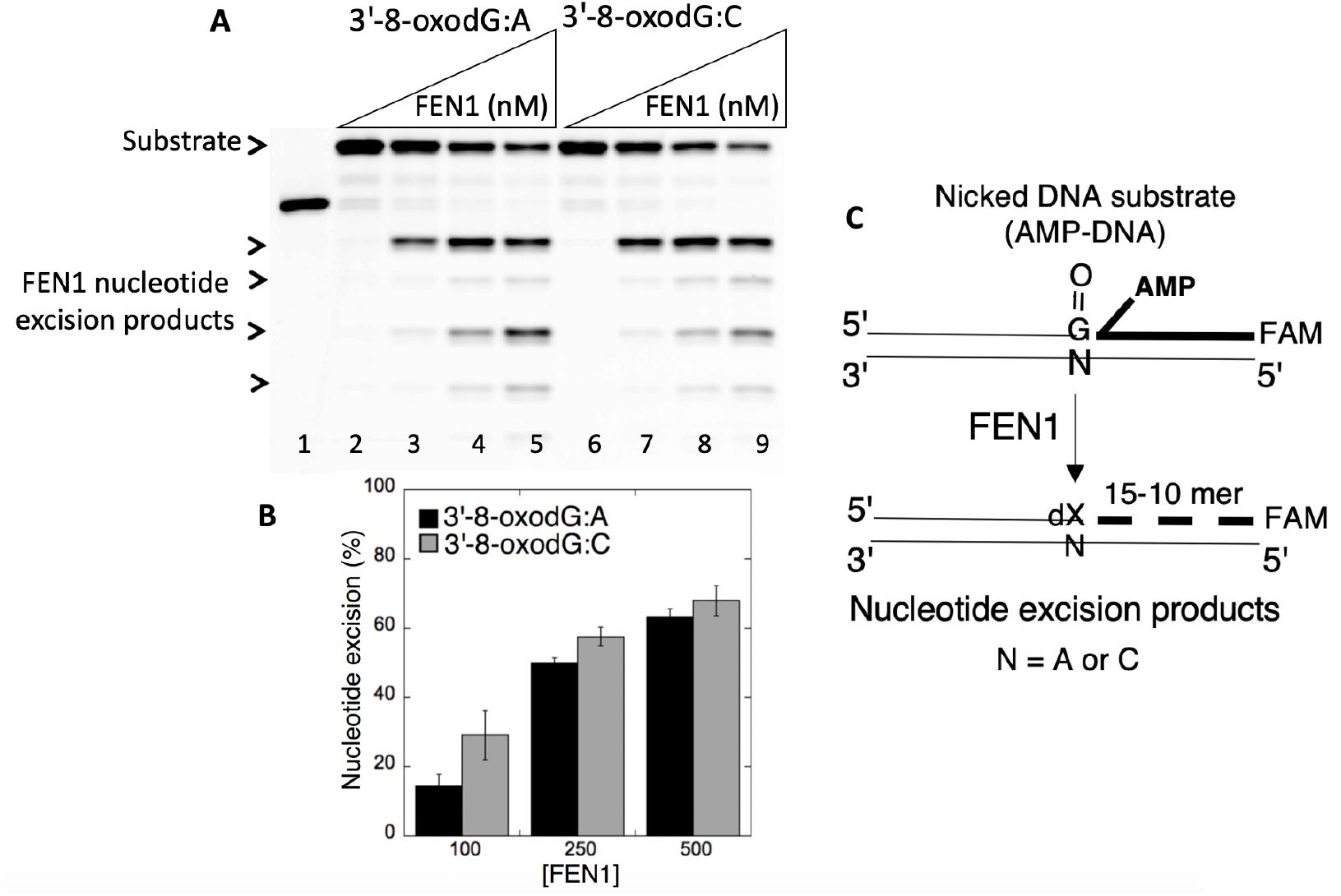
The nucleotide excisions by FEN1 from the ligation failure products with 3’-8-oxodG. (**A**) Lane 1 is a size marker that corresponds to the oligonucleotide without an AMP moiety. Lanes 2 and 6 are the negative enzyme controls of the nicked DNA substrates with 5’-AMP and 3’-8-oxodG opposite template base A and C, respectively. Lanes 3-5 and 7-9 show the products of nucleotide excisions from 3’-8-oxodG:A and 3’-8-oxodG:C substrates, respectively, obtained as a function of FEN 1 concentration. (**B**) The graph shows the time-dependent changes in the products of nucleotide excisions and the data are presented as the averages from three independent experiments ± SDs. (**C**) Illustrations of the nicked DNA substrate with 5’-AMP and 3’-8-oxodG and the products observed in the FEN1 reaction.

We then analyzed APTX and FEN1 activities for the repair intermediates with 5’-AMP and 3’-preinserted mismatches for all possible 12 non-canonical base pairs (Figures 13C and 14C). For example, for the nicked DNA substrates including template base A mismatches, we obtained the products of 5’-adenylate removal from DNA ends with 3’-preinserted base pairs dA:A, dC:A, and dG:A (Figure 13A, lanes 3-8, 9-14, and 15-20, respectively). Similarly, for the nicked DNA substrates including template base C mismatches, we obtained the products of 5’-adenylate removal and nucleotide excisions from DNA ends with 3’-preinserted base pairs dA:C, dC:C, and dT:C (Figure 14A, lanes 3-5, 7-9, and 11-13, respectively). Overall results demonstrated the products of 5’-AMP removal by APTX (Supplementary Figure 6) and the nucleotide excisions by FEN1 (Supplementary Figure 7) from all possible 12 mismatch-containing DNA substrates with no significant difference in the specificity for 3’-mismatch:template base combination (Figures 13B and 14B).

**Figure 13.**
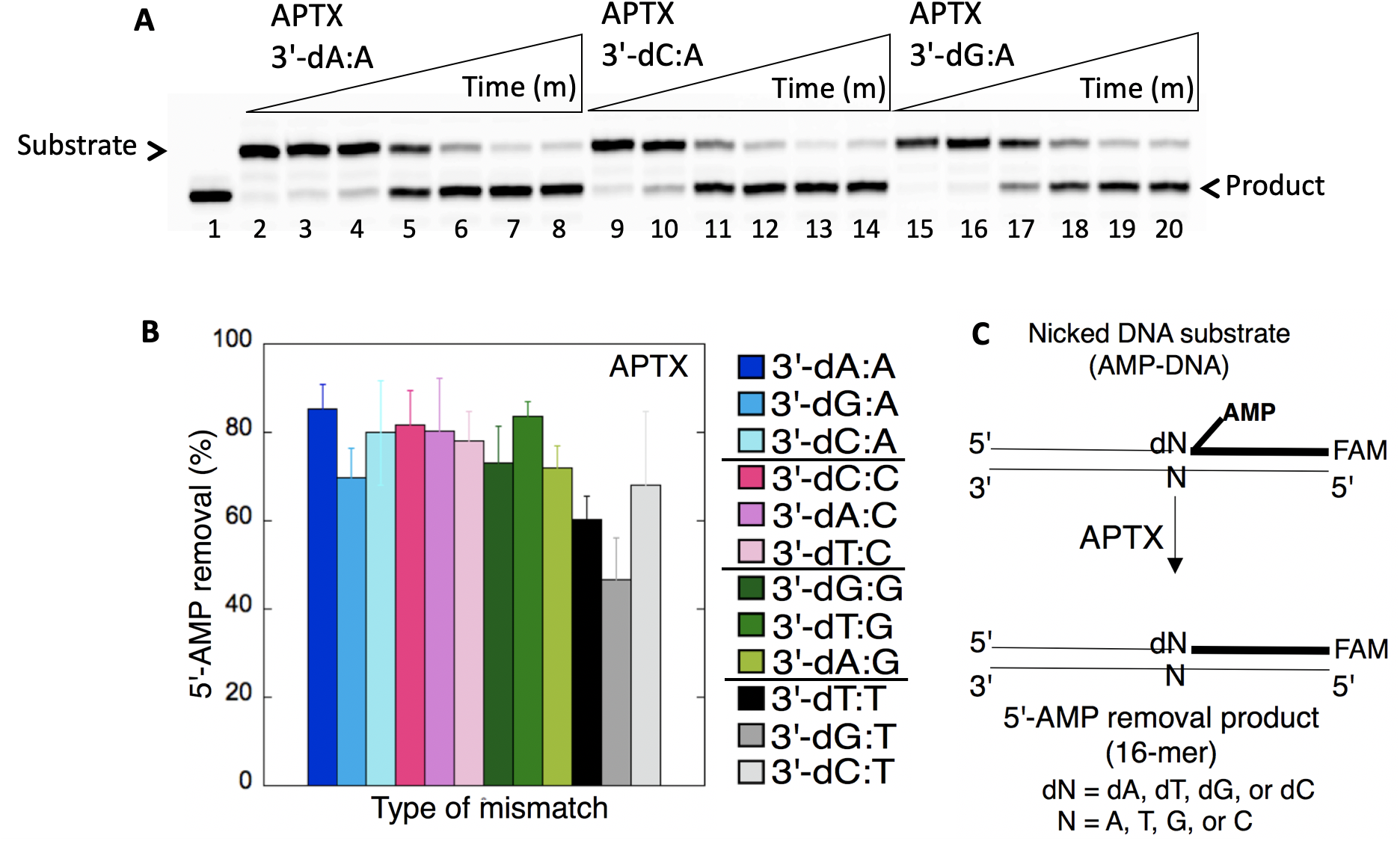
Removal of 5’-AMP by APTX from the ligation failure products with preinserted 3’-mismatches. (**A**) Lane 1 is a size marker that corresponds to the oligonucleotide without an AMP moiety and lane 2 is the negative enzyme control of the nicked DNA substrate with 5’-AMP and 3’-mismatched base. Lanes 3-8, 9-14, and 15-20 show the products of 5’-AMP removal from 3’-dA:A, 3’-dC:A, and 3’-dG:A mismatch-containing substrates, respectively, obtained at the time points 0.5, 1, 3, 5, 8, and 10 min. (**B**) The graph shows the comparisons in the products of 5’-AMP removal between all 12 non-canonical base pair mismatches. The data are presented as the averages from three independent experiments ± SDs. The gel images that show 5’-AMP removal from all other mismatches are presented in Supplementary Figure 6.

**Figure 14.**
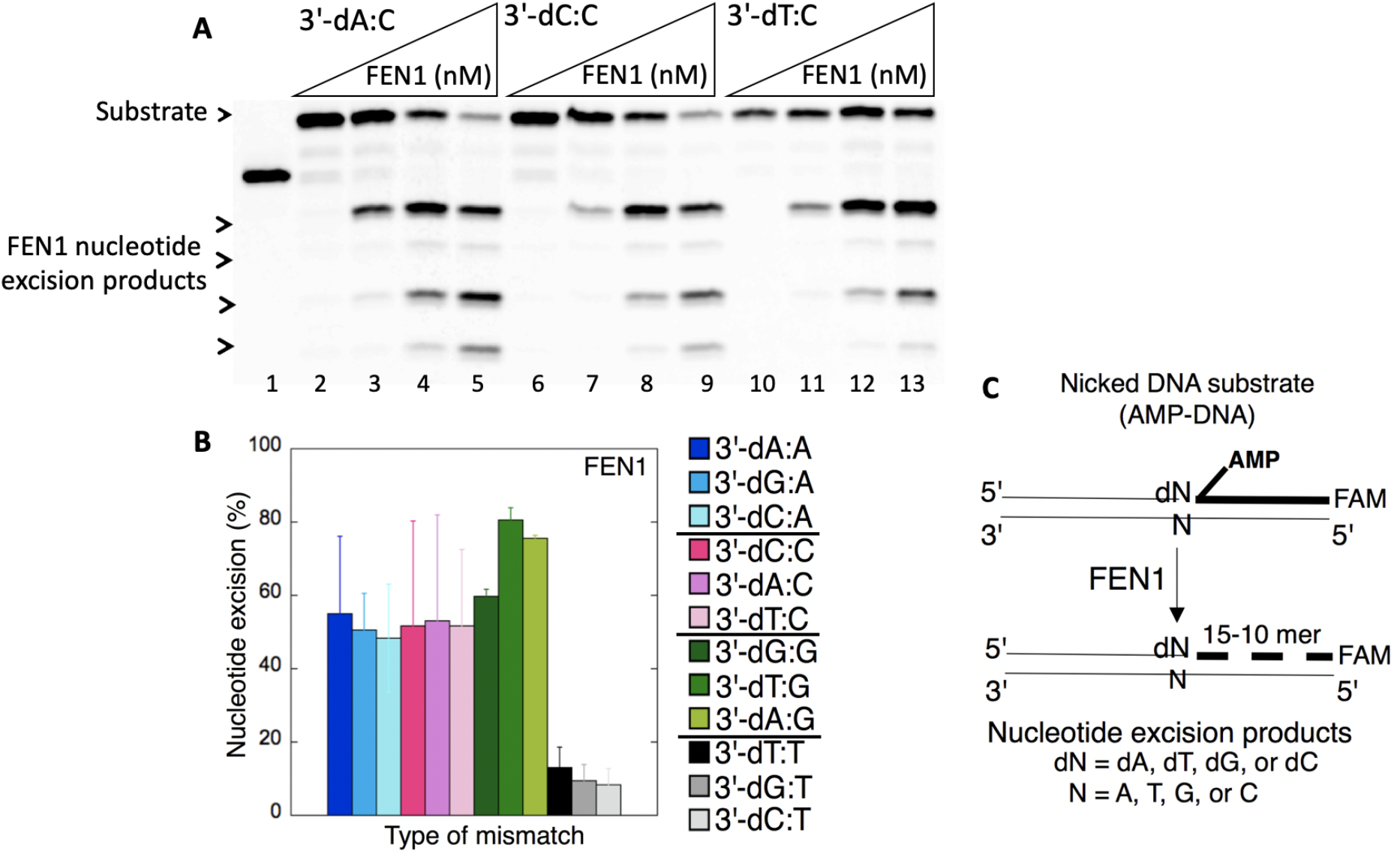
The nucleotide excisions by FEN1 from the ligation failure products with preinserted 3’-mismatches. (**A**) Lane 1 is a size marker that corresponds to the oligonucleotide without an AMP moiety. Lanes 2, 6, and 10 are the negative enzyme controls of the nicked DNA substrates with 5’-AMP and 3’-dA:C, 3’-dC:C, and 3’-dT:C, respectively. Lanes 3-5, 7-9, and 11-13 show the products of nucleotide excisions from 3’-dA:C, 3’-dC:C, and 3’-dT:C mismatch-containing substrates, respectively, obtained as a function of FEN1 concentration. (**B**) The graph shows the comparisons in the products of nucleotide excisions for all 12 non-canonical base pair mismatches. The data are presented as the averages from three independent experiments ± SDs. The gel images that show nucleotide excision products from all other mismatches are presented in Supplementary Figure 7.

## DISCUSSION

DNA ligases are essential enzymes for DNA replication and repair, and they join DNA strands to finalize each of these processes (1–4). DNA ligase I (LIG1) ligates the newly synthesized lagging strand during replication to complete Okazaki fragment maturation and is one of the key enzymes finalizing the BER pathway at the downstream steps after pol β gap filling or nucleotide insertion step (41). Recently published high resolution structures of LIG1 have identified the protein-DNA interface that is crucial for high fidelity (HiFi) DNA ligation (32). The X-ray structures of LIG1 bound to a nicked DNA duplex revealed the Mg^HiFi^ binding site that is coordinated by the LIG1 domains of Adenylation (AdD), DNA-binding (DBD), and the DNA phosphodiester backbone. This site is not found in other human DNA ligases III and IV and plays a vital role in identifying DNA ends during nick sealing and provides an additional checkpoint to hinder the abortive ligation. The mutations at the two conserved glutamic acid (E) residues (E346 and E592) to alanine (A) at the LIG1 active site (E346A/E592A) creates the ligase with lower fidelity due to an open cavity that accommodates and seals damaged DNA termini (32). However, the biochemical characterization of low fidelity LIG1 in terms of BER regulation is missing. In the present study, we investigated the pol β/LIG1 interplay in the execution of the different repair steps during substrate-product channeling at the downstream steps of BER pathway when LIG1 fidelity is perturbed by the E346A/E592A mutation (EE/AA LIG1). We also further investigated the role of the HiFi Mg^+2^ site of LIG1 on ligation fidelity of the nicked repair intermediates that mimic pol β oxidized or mismatch insertion products.

Our results revealed that the EE/AA LIG1 mutant is capable of mutagenic ligation by accommodating 8-oxodGMP inserted by pol β in a BER reaction. In contrast, wild-type LIG1 fails on the insertion product leading to the ligation failure intermediates with 5’-adenylate (AMP) block. Interestingly, the mutagenic ligation of the nicked pol β insertion product with 8-oxodGMP opposite A is much more efficient and faster than the ligation of pol β 8-oxodGTP insertion opposite C. This could be due to the distinct structural conformations of the primer terminus backbone with damaged termini at the ligase active site, which is stemming from dual coding potentials of 8-oxodG (*anti*- or *syn*-) as previously reported for pol β (25, 42).

On the other hand, our results showed that LIG1 with perturbed fidelity attempts to ligate the gap DNA substrate that pol β uses to insert dGTP mismatch opposite T in a BER reaction (referred as “self-ligation”) in contrast to the efficient ligation of the pol β repair product with G-T mismatch by the wild-type LIG1. This could result in potential single nucleotide deletion mutagenesis products and indicate a lack of effective substrate channeling. It is important to note that pol β active site with a G-T mismatch escapes mismatch discrimination through ionization of the wobble base pair (43,44). Moreover, Watson-Crick like G:T base pairing has been considered as an important source of base substitution errors that could be formed during DNA repair or replication, if left unrepaired, leads to transition or transversion point mutations in cancer (45–48). In our previous studies, we also demonstrated the difference in the substrate specificity of LIG1 when the nicked repair intermediate with a noncanonical base pair is handed from the pol β-mediated mismatch insertion step (16). Our overall findings suggest that the interplay between pol β and LIG1 in a multiprotein/DNA complex would be important for controlling the channeling of toxic and mutagenic DNA repair intermediates and the mutations that govern pol β and/or LIG1 fidelity could affect repair outcomes and faithful BER at the final steps (9). This could be biologically relevant in case of any cellular conditions related to the reduced fidelity of LIG1-mediated DNA repair or replication, which may contribute to the development of many diseases. For example, the reiteration of the mutations identified in the LIG1 gene of human patients who exhibit the symptoms of developmental abnormalities, immunodeficiency and lymphoma, in a mouse model has proven that LIG1 deficiency causes genetic instability and predisposition to cancer (49–53). Recently, we reported that the LIG1 variants associated with LIG1-deficiency disease exhibits different ligation fidelity for DNA polymerase-promoted mutagenesis products (14). Moreover, the studies have been suggested a pivotal role for pol β-mediated high-fidelity nucleotide insertion during gap filling step of BER pathway in the prevention of carcinogenesis (54).

In addition to our findings from coupled BER reactions including pol β and LIG1 as discussed above, we also obtained different ligation efficiencies in the ligation reactions including LIG1 alone for the nicked DNA substrates with 3’-8-oxodG opposite A or C vs G or T. Our results revealed the ligation failure of the repair intermediates containing preinserted 3’-8-oxodG when the damaged base is base paired with template G or T in contrast to the mutagenic ligation of DNA ends with 3’-8-oxodG opposite A or C by the LIG1 EE/AA mutant. Unfortunately, there is no LIG1 structure with this particular mismatch to be able to interpret our findings. According to the crystal structure of human LIG1 bound to adenylated DNA containing an 8-oxodG:A base-pair (PDB:6P0E), when the high fidelity residues E592 and E346 are mutated to alanine, Mg^+2^ binding site moves away from the active site leaving a cavity that can only be used for the damaged base (8-oxoG) in the downstream DNA strand (32). We suggest that the existence of G or T in the template DNA strand does not align the catalytic participants for optimal chemistry, prevent ligation of damaged termini, and require additional conformational adjustments at the LIG1 active site (EE/AA) to accommodate template base G or T while engaging the primer terminus with a damaged base to be ligated (Supplementary Figure 8). Similarly, the structural studies demonstrated that the pol β active site undergoes diverse mismatch-induced molecular adjustments and the extent of these conformational distortions in the pol β active site is dependent on the architecture of the mismatched template primer (55–59). We suggest that LIG1 poor active-site geometry observed for DNA ends containing 3’-8-oxodG opposite G or T could be an additional fidelity checkpoint that takes advantage of this poor geometry by deterring mutagenic ligation despite of the mutations at the HiFi sites. Similarly, our results demonstrated inefficient nick sealing of DNA ends with non-canonical base pairs 3’-dA:G or 3’-dG:A by the LIG1 EE/AA mutant. For DNA ligases from other sources, the study with *Tth* ligase from *Thermus thermophilus HB8* demonstrated that the base pair geometry is much more important than relative base pair stability and the ligase probes the hydrogen bond acceptors present in the minor groove to ensure the fidelity, which has also been proposed for DNA polymerases (60). Although it appears to be mechanistically similar to the incorporation of deoxynucleotide triphosphates by pol β, further structural studies are required to understand the mechanism by which the human LIG1 fidelity is accomplished in order to gain an insight into the mechanistic basis for discrimination against the range of substrates with aberrant base pair architecture.

As persistent DNA breaks are expected to be toxic, to block transcription and to be converted into double-strand breaks upon DNA replication, the formation and repair of the ligation failure intermediates are expected to be critical in cellular viability and genome stability (9, 10). In our previous studies, we reported the formation of 5’-AMP-containing BER intermediates after pol β 8-oxodGTP insertions *in vitro* and an increased cytotoxicity to oxidative stress inducing agent in pol β wild-type cells than pol β-null cells *in vivo* (13). The accumulation of adenylate intermediates in cells could be toxic leading to DNA strand break products with 8-oxodG or AMP lesions. Mutations in the APTX gene (*aptx*) are linked to the autosomal recessive neurodegenerative disorder Ataxia with oculomotor apraxia type 1 (AOA1) (33,37). APTX null cells fail to show a hypersensitivity phenotype to DNA damage-inducing agents and normal repair of base lesions and strand breaks have been reported in an APTX deficient mouse model (61–63). We previously reported the complementary role of FEN1 in case of deficiency in APTX enzymatic activity in the cell extracts from AOA1 patients (34–36). In the present study, our findings demonstrated the presence of the compensatory mechanism to deadenylate ligation failure products that could be formed because of pol β-promoted mutagenic nucleotide insertions and aberrant LIG1 fidelity. We showed the enzymatic activities of DNA-end processing proteins APTX and FEN1 that are indeed able to effectively remove 5’-AMP block from the repair intermediates containing with all possible 12 mismatched and oxidative damaged (8-oxodG)-containing DNA ends at 3’-terminus (*i.e*., pol β mismatch or oxidized base insertion products). On the other hand, in order to enable the repair to proceed, an additional 3’-end processing by a proofreading enzyme such as APE1 or mismatch repair protein (*i.e*., MSH2) is required for removal of damaged or mismatched base (64–66). Further structural and biochemical studies are required to understand how the repair enzymes (pol β, APE1, LIG1, APTX, and FEN1) function all together in a multi-protein/DNA complex to facilitate the faithful channeling of DNA repair intermediates. Moreover, in order to elucidate the potential contribution of crosstalk between these BER players and mismatch repair proteins as well as several proteins related to the DNA damage response (*i.e*., XRCC1 that interacts with pol β, APE1 and APTX), cell studies should be considered to gain further insight into the determinants of repair fidelity. In the future, it may be possible to develop small molecule inhibitors against the protein interacting partners of human LIG1 that could potentiate the effects of chemotherapeutic compounds and improve cancer treatment outcomes (9).

